# Lateral Asymmetry in Weekly Reproducibility of Resting State Amygdala Functional Connectivity

**DOI:** 10.1101/439372

**Authors:** Alina Tetereva, Vladislav Balaev, Sergey Kartashov, Vadim Ushakov, Alexey Ivanitsky, Olga Martynova

## Abstract

Abnormal functional connectivity of the amygdala with several other brain regions has been observed in patients with higher anxiety or post-traumatic stress disorder, both in a resting state and threatening conditions. However, findings on the specific connections of the amygdala might be varied due to temporal and individual fluctuations in the resting state functional connectivity (rsFC) of the amygdala and its lateral asymmetry, as well as possible variability in anxiety among healthy subjects. We studied reproducibility of rsFC data for the right and left amygdala, obtained by functional magnetic resonance imaging twice in a one-week interval in 20 healthy volunteers with low to moderate anxiety. We found resting-state amygdala network, which included not only areas involved in the emotion circuit, but regions of the default mode network (DMN) associated with memory and other brain areas involved in motor inhibition and emotion suppression. The amygdala network was stable in time and within subjects, but between-session reproducibility was asymmetrical for the right and left amygdala rsFC. The right amygdala had more significant connections with DMN regions and the right ventrolateral prefrontal cortex. The rsFC values of the right amygdala were more sustained across the week than the left amygdala rsFC. Our results support a hypothesis of functional lateralization of the amygdala. The left amygdala is more responsible for the conscious processing of threats, which may produce more variable rsFC; the right amygdala rsFC is more stable due to its greater engagement in continuous automatic evaluation of stimuli.

**Highlights:**

- Amygdala resting state network included areas of emotion circuit and motor control
- During rest amygdala was functionally connected with areas of default mode network
- Functional connectivity of the right amygdala was more sustained across the week
- Functional connections of amygdala network were more stable in the right hemisphere

## 1. Introduction

Several brain structures play a crucial role in learning emotionally significant stimuli. Some of these brain regions form the neuronal circuits of the fear network, as they are specifically involved in processing fear and anxiety. Task-based functional magnetic resonance imaging (fMRI) studies of fear conditioning in humans have found consistent, robust activation in response to conditioned stimuli in the amygdala, anterior cingulate cortex (ACC), anterior insula, and hippocampus (HIP), using both trace and delay conditioning paradigms in fMRI (Buchel et al., 1999; Pohlack et al., 2012; Andreatta et al., 2015) and magnetoencephalography (Balderston et al., 2014).

Animal models and translational studies have established the amygdala to be a central hub of neural circuits for threat detection and emotional learning (LeDoux, 2000; Davis, 2006; Akirav and Maroun, 2007; Etkin et al., 2009; Schumann et al., 2011). Due to its involvement in emotional learning, the amygdala has been reported to contribute significantly to social cognition (Brothers 1990, Adolphs 2001, Phelps 2006; Etkin et al., 2007; Bickart et al., 2014). Moreover, some studies have referred to the amygdala as a region of the default mode network (DMN) subsystems related to memory (Li et al., 2015) and self-referential cognition (Sheline et al., 2009). The amygdala is particularly associated with perceiving, evaluating, and learning possible threats from the environment, and consequently modulating behavioral responses. As part of the fear network, the amygdala is the most common region of interest in neuropathology studies, especially in anxiety and stress-related disorders. Task-based fMRI studies reported altered amygdala activation and amygdala-linked circuitry involving the medial prefrontal cortex (mPFC), insula, ACC, posterior cingulate cortex (PCC), and HIP (Nemeroff et al., 2006; Rauch et al., 2006; Etkin et al., 2007; Shin, 2010; Zhou et al., 2012) in post-traumatic stress disorder (PTSD). Elevated resting state functional connectivity (rsFC) of the amygdala with several other regions of the fear network has been also reported in PTSD patients (Bluhm et al., 2009; Daniels et al., 2010; Lanius et al., 2010; Liao et al., 2010; Dickie et al., 2011; Hayes et al., 2011; Rabinak et al., 2011; Yin et al., 2011; Zhou et al., 2012; Brown et al., 2014), as well in healthy groups that have undergone fear conditioning (Schultz et al., 2012), fear extinction, and fear reminding (Rauch et al., 2006). Anxiety level also influences amygdala connectivity. Kim et al. (2010) found a positive correlation between state anxiety scores and rsFC of the amygdala with mPFC. Baur et al. (2013) also reported a strong correlation of rsFC between the anterior insula and the basolateral amygdala with state anxiety scores.

Increased coupling of the amygdala with the ACC and the insula was replicated in many fear-conditioning studies (Etkin et al., 2007; Lanius et al., 2010; Shin 2010; Rabinak et al., 2011; Schultz et al., 2012; Brown et al., 2014). In addition, resting state research has reported altered FC of the amygdala with ventromedial prefrontal cortex (vmPFC), HIP, precuneus and PCC in PTSD patients (Dickie et al., 2011; Hayes et al., 2011; Zhou et al., 2012). However, a set of described brain areas, which is fictionally connected with the amygdala during rest or emotion-induced tasks, is volatile from one study to another. This inconsistency in data might be explained by temporal and individual fluctuations in rsFC of the amygdala and other brain regions as well as possible variability in anxiety between subjects. The current study aimed to examine the reproducibility of rsFC of the amygdala with other brain regions across a one-week period in a group of healthy volunteers with low variability in anxiety and depression scores. Moreover, we wanted to check the stability of rsFC of bilateral regions of the amygdala separately. Previously, asymmetrical functional activation, connectivity, and morphology of the left and right amygdala have been indicated in different patients’ groups (Hahn et al., 2011); however, it has rarely been examined in a healthy human population. In the first stage, we examined differences in the connectivity values of the left and right amygdala over a one-week interval using a seed-based approach. Then, in accordance with the seed-based data, we evaluated weekly changes in FC between several bilateral regions of interest (ROIs) that demonstrated significant correlation with a time-course of the blood oxygenation level dependent (BOLD) signal from lateral amygdala seeds, in the first, second, or both scanning sessions. In the final stage, we calculated intraclass correlation coefficients (ICC), defined by the interaction ratio of within- and between-subject variance (Shrout and Fleiss, 1979), in order to estimate test-retest reproducibility of the FC values obtained within a one-week interval.

## 2. Materials and methods

### 2.1. Participants

In total, 22 healthy volunteers participated in the study. Subjects were recruited via advertisement on mailing lists and notice boards at the Institute of Higher Nervous Activity and Neurophysiology of Russian Academy of Science (IHNA), Moscow, RF. Subjects were selected based on a preliminary survey for exclusion criteria and scores lower than 9 points on the Beck Depression Inventory (BDI). We used general exclusion criteria such as prior head trauma, any contraindications against MRI, medication affecting the central nervous system, history of neurologic or psychiatric disorders, consumption of drugs, excessive consumption of alcohol and nicotine, pregnancy, and age over 30. Of the 22 participants enrolled after the preliminary selection, two were excluded for excessive motion during scanning (more than 1.5 mm). All subjects were right-handed, according to self-report. All participants provided written informed consent to participate in the study, which was reviewed and approved by the ethics committee of IHNA, Moscow. The study procedure conforms to the Helsinki Declaration.

Overall, the data of 20 subjects (mean age 25.9 ± 4.27, 14 males and 6 females) were subjected to the further analysis (Table 1).

**Table 1.**
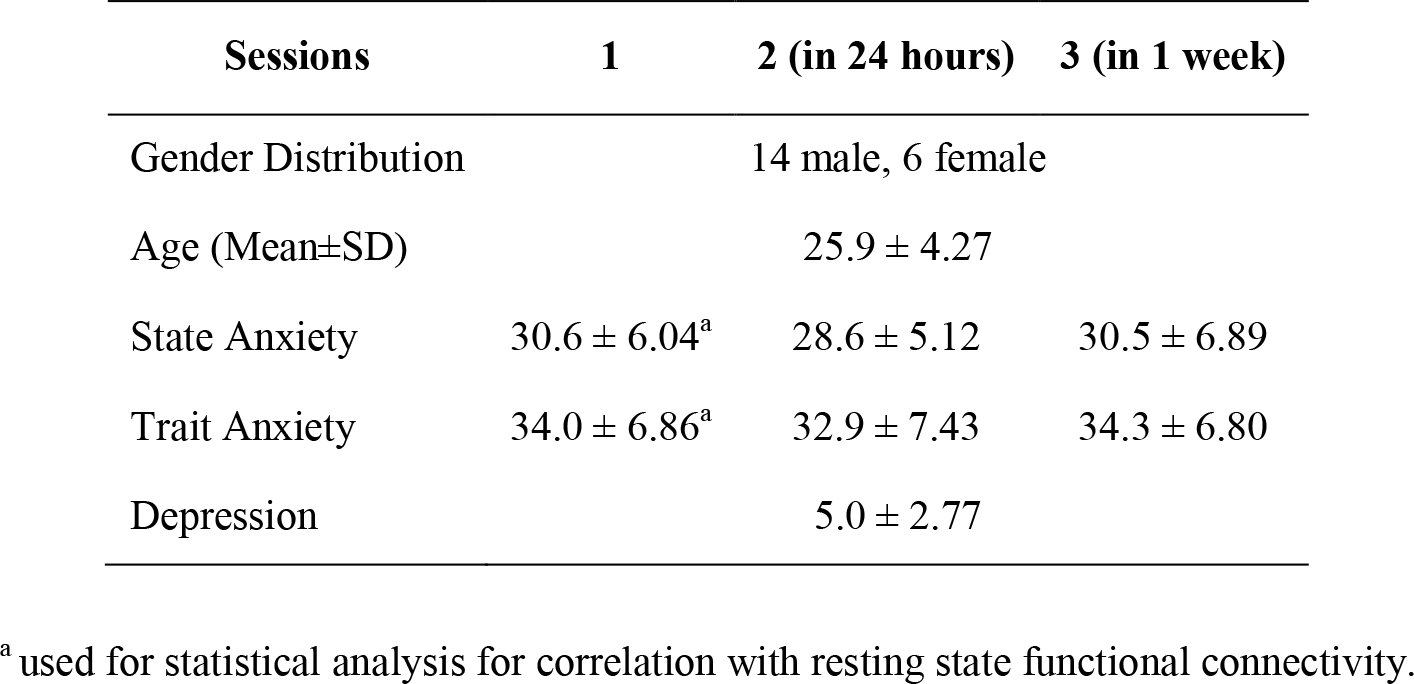
Demographic characteristics and state/trait anxiety scores of all participants.

### 2.2. Procedure

All participants underwent three scanning sessions: one session on the first day, a second session after 24 hours, and the third session seven days after the first one. Participants lay supine in a magnetic resonance imaging (MRI) scanner. They were instructed to remain calm, with their eyes closed, to remain awake and to avoid purposely thinking about anything. In addition, we stabilized the participants’ heads with the use of foam pads to diminish movement artifacts during MRI acquisition. In order to eliminate the possible influence of fear in the brain activity of participants who might have been experiencing MRI for the first time, or those who were afraid of or uncomfortable with the MRI procedure, we took data only from the second and third sessions for use in the test-retest reliability analysis and additional statistical comparison of the FC patterns between scanning sessions. For this reason, we refer hereinafter to the *second* scanning session as the *first analyzed session*, and to the *third* session as the *second analyzed session*. Before each scanning session, subjects completed the State-Trait Anxiety Inventory (STAI) (Spielberger et al., 1970).

### 2.3. Data acquisition

MRI data was collected at the National Research Center Kurchatov Institute (Moscow, RF) by a 3T scanner (Magnetom Verio, Siemens, Germany) equipped with a 32-channel head coil. For each subject, we acquired sagittal high-resolution T1-weighted rapid gradient-echo (anatomical) images using a T1 MP-RAGE sequence: TR 1470 ms, TE 1.76 ms, FA 9°, 176 slices with a slice thickness of 1 mm and a slice gap of 0.5 mm; and a field of view 320 mm with a matrix size of 320 × 320. We then collected functional images using a T2*-weighted echo planar imaging (EPI) sequence with a generalized autocalibrating partially parallel acquisition (GRAPPA) acceleration factor equal to 4 (Preibisch et al., 2008) and the following sequence parameters: TR 2000 ms, TE 20 ms, FA 90°, 42 slices with a slice thickness of 2 mm and a slice gap of 0.6 mm, a field of view (FoV) of 200 mm, and an acquisition matrix of 98 × 98. In order to reduce spatial distortion of EPI, we also acquired magnitude and phase images using a field map algorithm.

### 2.4. Preprocessing of MRI data

The MRI data was preprocessed in the Statistical Parametric Mapping version 8 (SPM8; Welcome Trust Centre for Neuroimaging, UK). Functional images were realigned with the mean functional image for motion correction. Magnitude and phase values from the field map images served for calculation of the voxel displacement map (VDM) and the subsequent mapping to the mean functional image. Next, each fMRI volume was unwrapped using VDM. After co-registration of the mean functional image with the anatomical image, all functional images were normalized into the standard template of Montreal Neurological Institute (MNI) space with a voxel size of 1.5 × 1.5 × 1.5 mm^3^ in two stages by the SPM8 New Segment tool: segmentation of anatomical images into grey and white matter, cerebrospinal fluid, bones, and air, followed by deformation of fMRI volumes and anatomical image using deformation fields. After the normalization procedure, we smoothed the fMRI images using a Gaussian kernel with full width at half maximum (FWHM) of 6 mm. Afterward, fMRI data were filtered by a fifth-order Butterworth band-pass filter with a frequency window from 0.01 Hz to 0.1 Hz.

During the final stage of preprocessing, we performed a regressing-out procedure in order to exclude motion patterns and signals derived from ventricles from the fMRI data (Fox et al., 2009). The mask for ventricles was created in WFU PickAtlas 3.0.4 (Maldjian et al, 2003) and then deformed to the individual space via the procedure described in the previous section. The ventricles’ time series and 6 motion parameters were extracted for each subject and used as regressors in the general linear model. The resulting regression residuals were used for further analysis of FC.

### 2.5. Seed-based analysis of amygdala’s functional connectivity

Masks of the amygdala were created for the left and right hemispheres in WFU PickAtlas 3.0.4 and co-registered with an individual anatomy image in the MNI space. They then underwent inverse deformation to the individual subject space for each participant. The last step included co-registration of the mask with the mean functional image. The mean BOLD signal was extracted for the left and right amygdala separately. Pearson correlation coefficients of the BOLD signal from the lateral amygdala seeds were calculated with a BOLD time series from every other voxel in the brain. Then the individual r statistics were normalized using a Fisher’s z transformation and subjected to a whole-brain one-sample t-test for both resting-state sessions (second day and in one week). Correction for false discovery rate (FDR) was applied to the results with a cluster threshold of more than 100 voxels (p < 0.001, T = 5.82). A two-sample t-test was also applied for comparison of resting-state correlation values between two scanning sessions.

### 2.6. Functional connectivity analysis between regions of interest

For functional connectivity analysis between regions of interest (ROI-FC), we selected brain areas that showed a significant increase of FC with either the right or left amygdala seeds in either of the two scanning sessions according to one-sample t-tests after FDR correction (p < 0.001, cluster threshold < 100). After that, 12 bilateral regions were chosen as ROIs for both hemispheres: amygdala (AMYG), insula (INS), HIP, parahippocampal gyrus (PHG), middle temporal gyrus (MTG), medial part of superior frontal gyrus (SFGmed), inferior frontal gyrus, pars orbitalis (IFGorb) and inferior frontal gyrus, pars triangularis (IFGtriang), caudate nucleus (CAUD), superior temporal pole - Brodmann area 38 (STP-BA 38), uncus (UNC) and precentral gyrus (preCG). All masks were extracted from the Automated Anatomical Labeling brain atlas (Tzourio-Mazoyer et al., 2002) except for UNC (from labels) and BA38 (from Brodmann areas) (Lancaster et al., 1997 & 2000; Maldjian et al., 2003 & 2004; Tziortzi et al., 2011).

Each mask was co-registered with an anatomy image in the MNI space and underwent inverse deformation to the individual subject space. The final step included co-registration of each mask with the mean functional image. The Pearson correlation of mean BOLD signals, extracted from each pair of ROIs, was calculated and normalized as in the seed-based analysis section. All FC values were corrected for multiple comparisons with family-wise error rate (FWE) (p < 0.05) and with the less-strict FDR at different p-thresholds (0.05 and 0.01).

We then calculated paired t-tests to estimate significant differences of ROI-FC between the sessions. The data from the selected ROI-FC values of two scanning sessions were subjected to analysis of variance (ANOVA) for repeated measures.

### 2.7. Test-retest reliability of resting state functional connectivity between scanning sessions

Test-retest reliability of ROI-FC and sbFC was estimated by calculating the ICC (Shrout and Fleiss, 1979). We used a case “1-k” designed to evaluate the degree of absolute agreement of measurements that are averages of k-independent measurements on randomly selected objects. Mean ICC values were calculated inside masks of resting state networks from the CONN brain connectivity toolbox (Whitfield-Gabrieli and Nieto-Castanon, 2012) and subjected to one-way ANOVA. The obtained ICC values were also corrected for multiple comparisons with FWE (p < 0.05) and with the less-strict FDR at different p-thresholds (0.05, 0.01, and 0.001).

### 2.8. Correlation of amygdala connectivity with anxiety and depression scores

In order to increase the sensitivity of our reproducibility analysis to individual differences in FC and psychological scores, we tested a relationship between individual values of the self-report scores for state/trait anxiety, and depression with FC values for both scanning sessions, using normalized values in a Pearson correlation analysis. Following this, we ran separate multiple regressions in SPM8 with testing scores of state/trait anxiety and depression as regressors and amygdala connectivity as the dependent variable. We tested a relationship between amygdala FC for the first and second sessions. State/trait anxiety and depression scores were taken only on the first day as we did not find any difference between days using a paired sample t-test. Additionally, we assumed that only the first test results were more valid. Participants answered self-report questionnaires more objectively the first time they took the test than on repeated tests at short time intervals, as in our study. In repeated tests, participants may provide rehearsed answers because they usually remember their answers from the previous testing session.

## 3. Results

### 3.1. Anxiety scores did not change in one week

All selected subjects had self-report scores for anxiety and depression within the normal range (Table 1). There were no significant differences in STAI and BDI scores among the three sessions. The group of participants had low or moderate scores of state anxiety (63% low and 37% moderate, ranging from 22–43 before the first scanning session). Trait anxiety was mostly at a moderate level (31% low, 56% moderate, and 13% high rates, ranging from 24–49 before the first scanning session). All participants had low (< 9) BDI scores. Furthermore, we did not observe significant differences in the STAI and BDI scores between males and females.

### 3.2. Amygdala connectivity with other brain regions for two sessions at a one-week interval

A whole-brain two-tailed one-sample t-test performed for two sessions separately showed different brain regions in which the BOLD fluctuation time series correlated significantly with the left or right amygdala (Table 2). Spatial maps demonstrated both lateral and temporal differences in the correlation of BOLD signal in the brain regions with the mean signal from the left and right amygdala for two scanning sessions (Figure 1). In the first session, we observed significant positive FC of the left amygdala with three distinct areas during 10 min of the resting state condition: right amygdala, left MTG, and right Caud. The right amygdala showed significantly stable and positive connectivity with seven areas: right BA38, left Hip, right IFGorb, right Ins, right MTG, right IFGtriang, and left SFGmed.

**Table 2.**
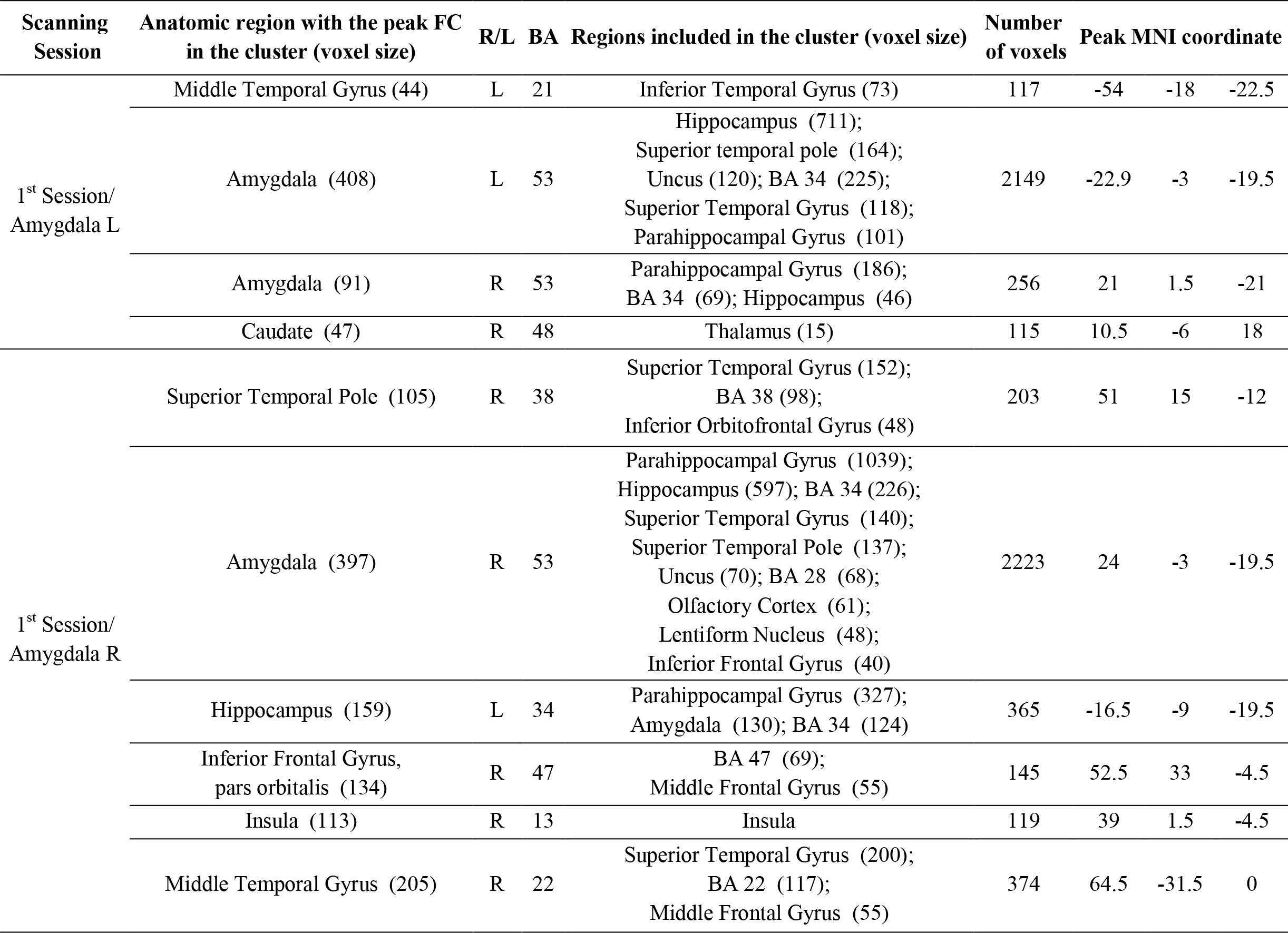
FC of the left and right amygdala with other brain regions according to separate one-sample t-tests for the first and second sessions in a one-week period (FDR corrected, p < 0.001, T = 5.82, cluster threshold > 100). R-in the right, L- in the left hemisphere; BA-Brodmann area.

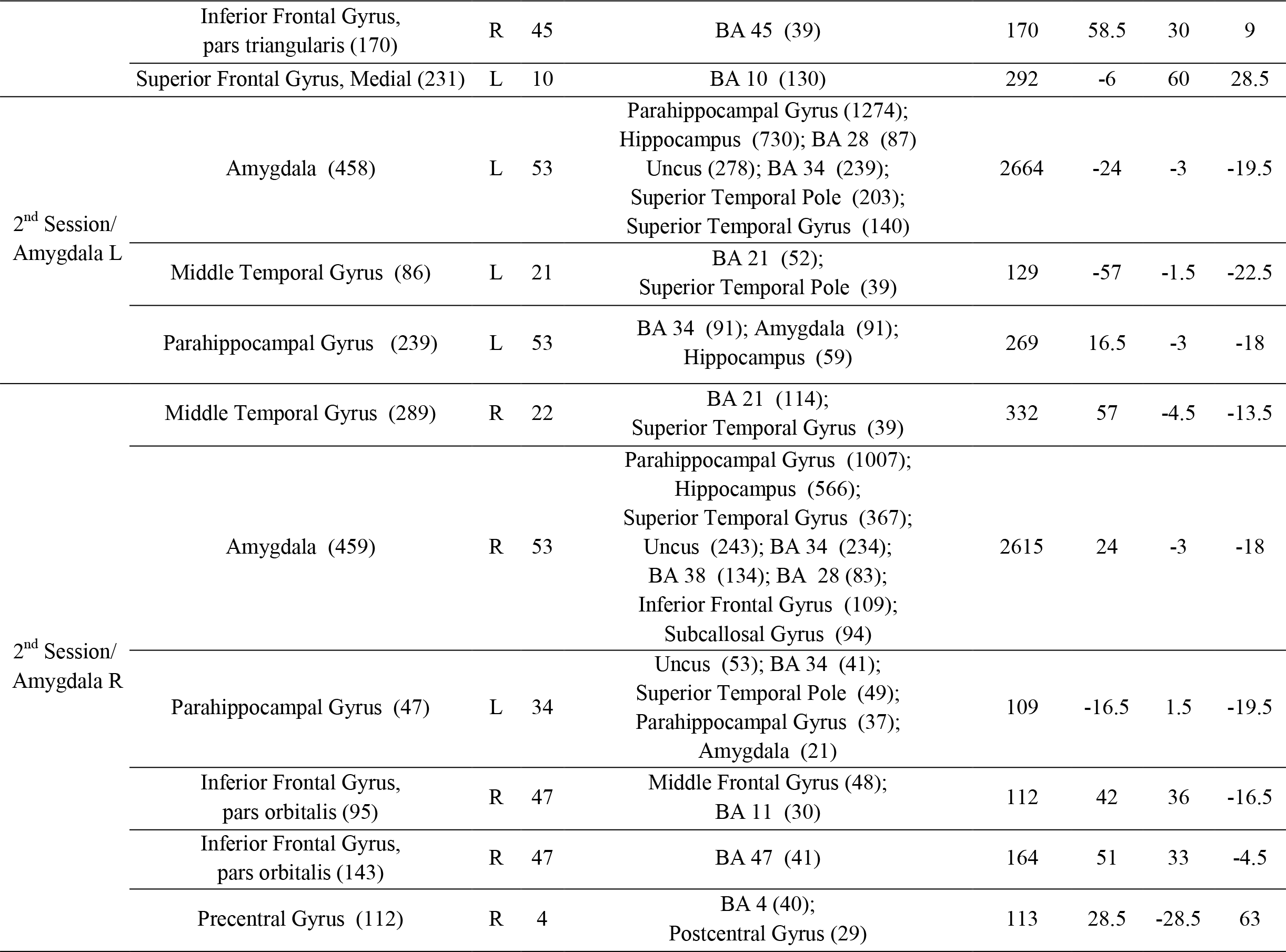

**Figure 1.**
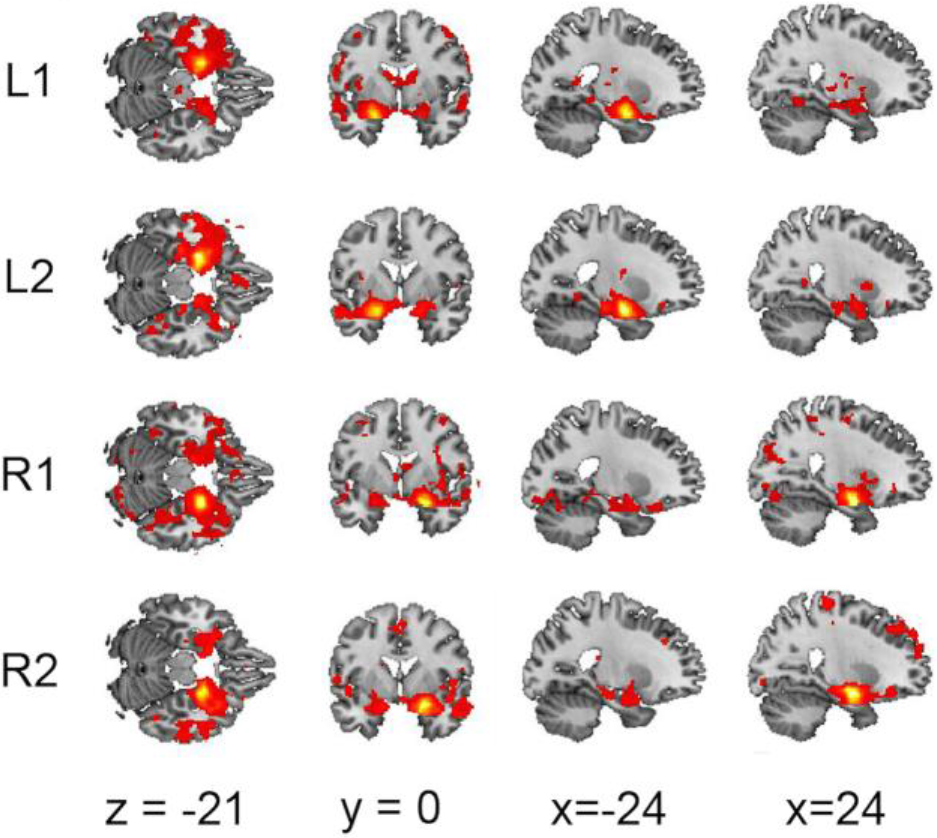
Significant positive correlations of functional connectivity for (L) the left and (R) right amygdala with other brain regions overlapped for two scanning sessions (1 – first session and 2 – in 1 week).

For the second session data (at a one-week interval), we found a significant positive correlation of the left amygdala with two brain areas: the left MTG and the left PHG. The right amygdala had five areas with highly stable FC: the right MTG, the left PHG, two clusters in the right IFGorb, and the right preCG. Thus, both the right and left amygdala correlated significantly with the left PHG in the third session.

Most areas showing the highest connectivity with the amygdala coincided for both sessions, especially MTG. Moreover, both lateral parts of the amygdala expressed a stable FC with ipsilateral MTG in the two studied sessions. The localization of the maximal value of FC within a cluster varied between sessions, but areas included in the clusters remained the same. These areas of the significant positive FC with the amygdala were either merged or split into smaller ones, depending on the session. (Table 2, Figure 1). Respectively, we found no significant effect of the session using a 2-sample t-test for direct comparison of the amygdala rsFC values. We did not observe even an uncorrected significant difference in the spatial distribution of FC values for either left or right amygdala connectivity.

### 3.3. ROI-FC changes and reproducibility of resting state functional connectivity between two scanning sessions at a one-week interval

Next, we analyzed pair-wise FC values of the brain areas that showed the increased connectivity with either the left or right amygdala in at least one of the two sessions. Figure 2 illustrates correlation matrices for 24 bilateral ROIs, including the left and right amygdala. Across 276 paired correlation coefficients, only 69 passed through FDR correction for multiple comparisons (p < 0.05) of the ICC reproducibility index computed for the two sessions. Of these, 21 pairs had some level of Pearson correlation r > 0.5 (Table 3), with steady bilateral coupling between Caud, Ins, and SFGmed. We observed a predominance of the stable connections in the right hemisphere (Figure 3). The majority of reproducible FC values were obtained for the right IFGorb and IFGtriang, with other ROIs in both hemispheres. The bilateral parts of preCG had an almost equal number of stable FC values, but with dominance of the right preCG. Additionally, we examined within-subject reliability of the ROI-FC values for each session separately, using a more rigorous FWE threshold. Some instability of FC across one week was found: 53 FC values were significant for the first session but only 44 ROI-FC values passed the significance threshold for the second session (Table S.1). FC between bilateral areas of SFGmed and Ins were the highest for both sessions.

**Table 3.** Regions of interest (ROI) with the most stable values of functional connectivity (z-scores of Pearson correlation coefficients) for the first and second sessions across a week, according to interclass correlation coefficients (ICC) (R-in the right, L- in the left hemisphere).

| Regions of interest | ICC<sup>a</sup> | z-score for 1st Session<sup>b</sup> | z-score for 2nd Session<sup>b</sup> | Mean z-score for 2 sessions<sup>c</sup> |
| --- | --- | --- | --- | --- |
| SFGmed L - SFGmed R | 0.67 | 1.25 | 1.26 | 1.26 |
| INS L - INS R | 0.70 | 0.66 | 0.81 | 0.74 |
| IFGorb R - SFGmed R | 0.75 | 0.69 | 0.67 | 0.68 |
| IFGorb R - MTG R | 0.74 | 0.66 | 0.66 | 0.66 |
| Caud L - Caud R | 0.65 | 0.62 | 0.68 | 0.65 |
| IFGorb R - SFGmed L | 0.67 | 0.65 | 0.62 | 0.64 |
| INS L - preCG L | 0.82 | 0.59 | 0.62 | 0.61 |
| SFGmed L - preCG L | 0.72 | 0.59 | 0.62 | 0.61 |
| IFGtriang R - preCG R | 0.78 | 0.61 | 0.60 | 0.60 |
| INS R - preCG R | 0.74 | 0.58 | 0.59 | 0.59 |
| IFGtriang R - INS R | 0.74 | 0.56 | 0.59 | 0.58 |
| IFGtriang R - preCG L | 0.66 | 0.56 | 0.57 | 0.57 |
| IFGorb R - MTG L | 0.65 | 0.57 | 0.55 | 0.56 |
| INS L - preCG R | 0.82 | 0.52 | 0.59 | 0.56 |
| SFGmed L - preCG R | 0.68 | 0.52 | 0.59 | 0.56 |
| BA38 R - MTG R | 0.73 | 0.56 | 0.55 | 0.56 |
| IFGtriang R - INS L | 0.74 | 0.51 | 0.60 | 0.55 |
| SFGmed R - preCG L | 0.71 | 0.51 | 0.59 | 0.55 |
| BA38 R - IFGorb R | 0.72 | 0.51 | 0.54 | 0.53 |
| SFGmed R - preCG R | 0.68 | 0.47 | 0.57 | 0.52 |
| IFGorb R - preCG R | 0.76 | 0.53 | 0.50 | 0.52 |

<sup>a</sup> $p < 0.05$ after FDR correction;
<sup>b</sup> $p < 0.05$ after FWE correction;
<sup>c</sup> Table values are sorted according to the decreasing mean z-scores for both sessions and limited to $z > 0.5$ .

**Figure 2.**
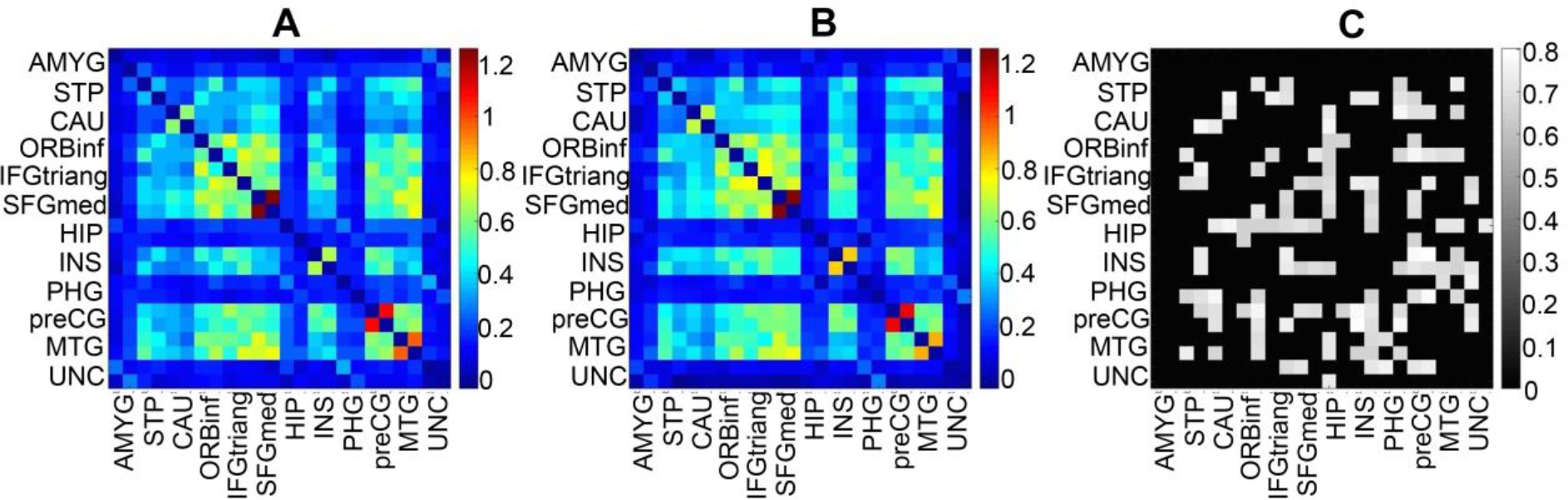
Cross-correlation maps of functional connectivity between regions of interest for two scanning sessions (A – first session, B – after a week). Each first row or column for region corresponds values in the right hemisphere, each second – in the left hemisphere. Significant differences are marked by points. Matrix C shows the most stable FC across a week with ICC > 0.4, FDR = 0.05.

**Figure 3.**
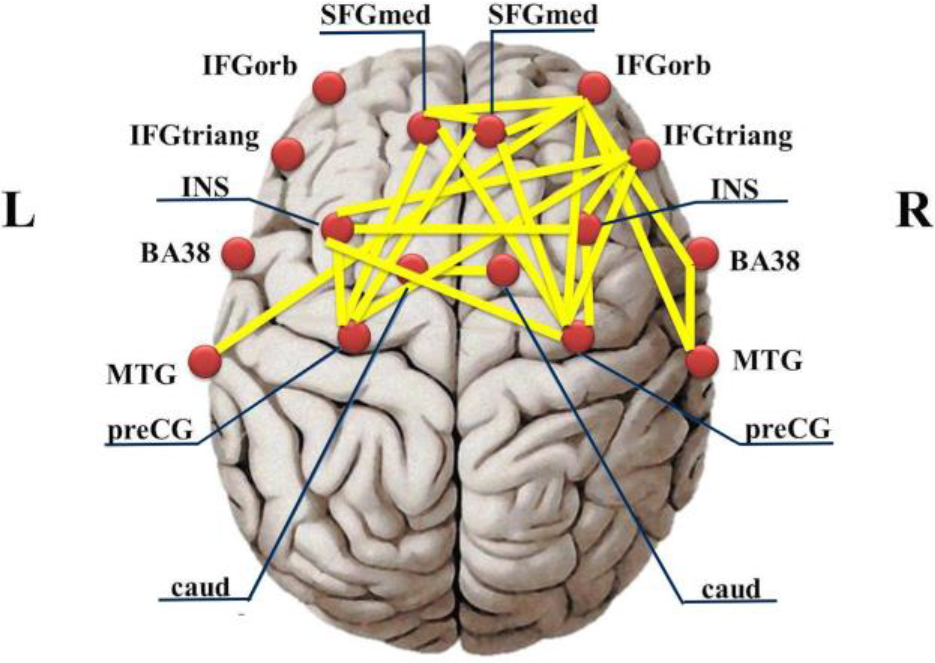
Stable functional connectivity between regions of interest with interclass correlation coefficients ICC < 0.7 (p < 0.05, FDR corrected) for two scanning sessions in a one-week interval.

FC of the amygdala with other ROIs was at a rather low level of Pearson correlation (0.1 < r < 0.23). Moreover, only 15 values of amygdala FC with other brain regions had reliable reproducibility (ICC > 0.4). The right amygdala occurred in 11 pairs; as for the left amygdala, we observed only 4 stable pairs (Table 4). None of these FC values of the amygdala passed through FDR correction (p < 0.05) except the FC of the right amygdala with the right preCG.

**Table 4.** Z-scores of Pearson correlation coefficients and interclass correlation coefficients (ICC) only for the left and right amygdala’ connections with regions of interest (R-in the right, L- in the left hemisphere).

| Regions of interest | ICC | z-score for<br>1st Session | z-score for<br>2nd Session | Mean z-score<br>for 2 sessions<sup>a</sup> |
| --- | --- | --- | --- | --- |
| AMY R - MTG R | 0.43 | 0.23** | 0.23** | 0.23 |
| AMY R - IFGorb R | 0.56 | 0.20** | 0.21** | 0.21 |
| AMY R - INS R | 0.49 | 0.17 | 0.19** | 0.18 |
| AMY R - preCG R | 0.66* | 0.19** | 0.17** | 0.18 |
| AMY L - HIP L | 0.59 | 0.20** | 0.16** | 0.18 |
| AMY R - preCG L | 0.57 | 0.18** | 0.17** | 0.18 |
| AMY R - HIP R | 0.51 | 0.16 | 0.14 | 0.15 |
| AMY R - PHG R | 0.44 | 0.17** | 0.12** | 0.14 |
| AMY R - INS L | 0.54 | 0.15** | 0.12 | 0.14 |
| AMY R - SFGmed R | 0.43 | 0.14 | 0.12 | 0.13 |
| AMY R - IFGtriang L | 0.49 | 0.14 | 0.10 | 0.12 |
| AMY L - BA38 L | 0.40 | 0.10 | 0.14** | 0.12 |
| AMY L - IFGorb L | 0.47 | 0.11 | 0.13 | 0.12 |
| AMY R - Caud R | 0.50 | 0.09 | 0.12 | 0.11 |
| AMY L - MTG L | 0.46 | 0.08 | 0.12 | 0.10 |

<sup>a</sup> Table values are sorted according to the decreasing mean z-scores for both sessions and limited to $z > 0.1$ ;
\* $p < 0.05$ after FDR correction;
\*\* $p < 0.05$ after FWE correction.

Among these 15 FC values (ICC > 0.4, r > 0.1), 7 were significant in terms of within-subject reliability (pFWE < 0.05) for the first session and 8 FC values for the second session. Notably, reliable but low FC of the right amygdala with the left insula was observed for the first session; for the second session, we found reliable FC of the right amygdala with right insula and left amygdala with left BA38 (Table 4).

Thus, our results repeatedly demonstrate lateral asymmetry in the amygdala FC in terms of both within-subject reliability and temporal reproducibility of data. In the resting-state condition, activity of the right amygdala demonstrates more persistent connections with other brain regions.

### 3.4. Significant correlations of FC with anxiety scores

The values of ROI-FC did not show a correlation with BDI scores. We found a significant positive correlation of trait anxiety scores only with the FC of the left Caud left and the right preCG (r = 0.58, p = 0.007 and r = 0.62, p = 0.003 for the first and second session correspondingly) and the FC of the left Caud left with the right IFGorb, for both sessions (r = 0.48, p = 0.032 and r = 0.45, p = 0.047 for the first and second session correspondingly).

## 4. Discussion

In the current study, we demonstrated that the resting-state FC of the amygdala displays features of lateral asymmetry in healthy subjects with normal anxiety scores. Specifically, we observed more brain areas showing stable connectivity with the right amygdala than with the left within subjects, for two scanning sessions separated by one week. This finding was supported by data of both the FC of the lateral amygdala seeds and the ROI-FC.

First, we observed significant lateral differences in the correlation of brain regions with the mean signal from the left and right amygdala for both scanning sessions. Although a distribution of the areas correlated with lateral amygdala seeds differed for the two sessions, the most areas showing within-subject reproducibility of FC values with the amygdala in each hemisphere overlapped for both sessions. While the left amygdala FC significantly coupled with ipsilateral MTG and PHG, contralateral Caud, and amygdala, the right amygdala showed a positive correlation of BOLD fluctuations with more brain regions, including not only areas involved in the limbic system and emotion processing (Caud, Hip, Ins, MTG, STP-BA38, and PHG), but also cortical regions associated with higher brain functions, such as the dorsomedial prefrontal cortex (SFGmed), right homologues of the language areas IFGorb and IFGtriang, and the motor cortex (preCG). Functional and anatomical connections of the amygdala with the limbic system and memory (Hip, Ins, and MTG) have previously been discussed in terms of emotional learning (LaBar et al., 2006). At the same time, Hip with PHG, MTG, STP, and SFGmed were consistently reported as DMN nodes (Raichle et al., 2001; Bukner et al., 2008; Laird et al., 2009; Andrews-Hanna et al., 2010). Our results show that the right amygdala interacts with areas of the DMN more than the left does during the resting-state condition. Remarkably, we did not observe interactions of the amygdala with vmPFC, OFC, or ACC, or with other neocortical areas having functional and anatomical connections with the amygdala and association with emotion processing (Carmichael & Price 1995; Bickart et al., 2014). On the contrary, in our study the amygdala demonstrated stable FC with other regions of the DMN, such as Hip, PHG, and MTG; these are strongly related to memory functions (Squire et al., 2015). Together with the amygdala, these structures are also anatomically binded and involved in declarative and episodic memory (Squire, 2009). Activity of the medial temporal lobe subsystem of the DMN has been shown to increase in memory-related tasks (Andrews-Hanna et al., 2010), effective connectivity between regions of the DMN, including Hip, PHG, and amygdala (and PCC), and are also correlated in both the resting state and active tasks with memory performance in healthy subjects (Li et al., 2015). Thus, our results indicated that in the resting-state condition, without any emotional stimulation or active task, the amygdala is more involved in memory than in emotional circuits.

In our study, the right amygdala additionally expressed persistent connectivity with ipsilateral areas in IFGorb (BA 47) and IFGtriang (BA 45). These areas of IFG in the right hemisphere are mostly known as key parts of the brain’s language system (Düzel et al., 2001; Wildgruber et al., 2005), but they have also been reported to contribute to motor inhibition (Marsh et al., 2006; Völlm et al., 2006), and decision making (involving conflict and reward) (Rogers et al., 1999). Cha et al. (2016) highlighted a role of the left IFG in evaluation of stimulus meaning. During a fear generalization task, the FC of the left IFG with the vmPFC and the amygdala correlated significantly with severity of anxiety; however, no significant group differences between clinically anxious and healthy human subjects were found for the right IFG (Cha et al., 2016). In the meta-analysis of vlPFC, Levy and Wagner (2011) reported involvement of the IFGtriang in motor inhibition (Levy and Wagner, 2011). Additionally, the increased activation of the right IFG was mentioned in tasks requiring suppression of emotions (Vanderhasselt et al., 2013). In our study, increased connectivity of the right amygdala with vlPFC (IFGtraing and IFGorb) may indicate a possible role of the amygdala in motor inhibition, in a condition of constrained movement during MRI scanning. Moreover, the observed connectivity of the right amygdala with the preCG, or the primary motor cortex, which is responsible for motor control (Porro et al., 1996), supports the latter assumption about motor inhibition and/or motor imagination during a resting state. The left amygdala also showed constant connectivity during the first scanning session with Caud. The caudate nuclei of the dorsal striatum are known to be involved in sensorimotor and inhibitory control, but they also play an important role in the emotional learning and reward system (Grahn et al., 2008; Cha et al., 2016). Consequently, our findings of a steady coupling of the amygdala with brain areas of the motor control network suggest that both lateral parts of the amygdala also take part in inhibitory control of motor functions during a resting-state condition.

Furthermore, we performed ROI-FC analysis for the selected brain areas, which showed a significant increase of FC with either the right or left amygdala in either of the two scanning sessions. We found a predominance of the highest (r > 0.5, p < 0.001) and reproducible connections between ROIs in the right hemisphere. Apart from bilateral connections between SFGmed, Ins, and Caud, the majority of reproducible FC values were observed for IFGorb, IFGtriang, and preCG in the right hemisphere (Figure 3).

As the right hemisphere FC values of IFGorb, IFGtriang, and preCG were the most reproducible in time and across healthy subjects, it is natural to assume that these stable connections reflect cognitive control of movement inhibition in the resting-state condition. Since these areas had a substantial functional connection with the right amygdala, either in the first or the second session according to seed-based FC analysis, our findings imply that the right amygdala plays a certain role in cognitive control of movement based on its continuous involvement in evaluation of external stimuli.

According to our data, the amygdala showed a low coupling with other brain regions in the resting state in healthy subjects without induced fear. However, amygdala FC values were reproducible for several ROIs (ICC > 0.04); this was greater for the right amygdala than for the left.

Lateral asymmetry of the amygdala has previously been shown in many other studies. For example, Pedraza et al. (2004) reported that the structural volume of the right amygdala was larger than the left in healthy participants. However, a functional asymmetry of the amygdala and its connections has been discussed primarily in neuroimaging studies, either with induced emotions or in the comparison of FC in healthy and patient groups. Therefore, in the meta-analysis of the fMRI of emotions in healthy participants, Baas et al. (2004) concluded that activity of the left amygdala prevails in neuroimaging studies of emotional processing. Nevertheless, data on functional asymmetry of the amygdala in anxiety and stress-related disorders are controversial. For example, studies of FC in PTSD patients mainly reported the increased FC of the right amygdala with limbic brain areas (Bluhm et al., 2009; Lanius et al., 2010; Rabinak et al., 2011; Zhou et al., 2012). Motzkin et al. (2015) also observed significantly greater activation of the right amygdala in patients with bilateral damage of the vmPFC in response to aversive images, compared to the control group. However, some studies showed a stronger connectivity of the left amygdala with areas involved in emotional-motivational circuits in the resting state of PTSD groups (Dickie et al., 2011; Brown et al., 2014). Moreover, the meta-analysis study of Shin et al. (2010) of stress and anxiety disorders highlighted the increased activity of the left amygdala as an indicator of these diseases.

Functional lateralization of the amygdala may also depend on the subject’s awareness of environmental stimuli. The left amygdala expressed greater activation when participants knew and anticipated the unpleasant stimuli, while the right amygdala was more active when subjects were not aware of the presented stimuli (Morris et al., 1998; Phelps et al., 2001). This observation was further supported by data from meta-analyses, showing the greater involvement of the right amygdala in automatic processing of environmental stimuli, and on the other hand, greater engagement of the left amygdala in continuous evaluation of potential threats (Phan et al., 2002; Costafreda et al., 2008). Our findings may also support functional lateralization of the amygdala, depending on conscious processing of stimuli. First, we showed that the functional network of the amygdala during rest included not only areas of neural circuits of emotions but also brain regions of the DMN associated with resting-state cognition and memory (Raichle et al., 2001; Sheline et al., 2009). Second, this network had lateral asymmetry, as the right amygdala had more stable connections that were replicated in one week than the left amygdala. Thus, we may assume that the functional network of the right amygdala presented here is more permanent because it continuously maintains automatic/subconscious processing of stimuli. The left amygdala network, however, is more variable due to its attended/conscious evaluation of negative emotional stimuli. The impairment of the latter process might partially explain the lateral asymmetry of amygdala activation in various psychiatric disorders (such as obsessive-compulsive disorder or social anxiety disorder), where it has been shown that only the left amygdala connectivity differed in patients as compared with healthy controls (Hahn et al., 2011; Prater et al., 2013; Rus et al., 2017). Since we acquired resting-state fMRI data in participants with low or moderated anxiety twice in one week, the connectivity of the left amygdala might fluctuate substantially without experimental induction of negative emotions. Further research with subconscious fear conditioning and fear extinction may shed light on the lateral asymmetry of the amygdala’s functional connections.

In addition, we found significant correlations of trait anxiety scores between the FC of the left Caud with the right IFGorb and the right preCG. Apart from their established roles in motor inhibition, these are also involved in suppressing emotion, emotional learning, and reward systems (Vanderhasselt et al., 2013; Cha et al., 2016). However, we did not observe any significant correlations of STAI scores with the amygdala’s FC. Previous studies have also shown some inconsistency in results for correlation of FC with anxiety scores. Therefore, Baur et al. (2013) reported the left-lateralized FC of the amygdala, which correlated significantly with the anxiety level in healthy participants. In contrast, Kim et al. (2010) observed that FC between the right amygdala and the dorsomedial prefrontal cortex was positively correlated with scores of state anxiety. Hayes found a negative correlation between hippocampal activity and Clinician-Administered PTSD Scale scores. Liao et al. (2010) also reported a correlation between the Liebowitz Social Anxiety Scale scores with the amygdala’s effective connectivity network. In our case, the correlation results may be insufficient due to small sample size (20 participants) and low dispersion of low-to-moderate STAI scores between subjects. However, we aimed to investigate the FC of the amygdala during rest in healthy participants and used STAI assessment as additional criteria for selection of participants. More rigorous methods will be required in order to find significant connections between low-moderated anxiety and the amygdala’s FC at rest. The other limitation of the study related to a possible effect of weekly conscious experience of the participants on the amygdala’s FC fluctuations. Although we estimated the participants’ subjective stress by psychological questionnaires, we cannot exclude the possibility that stressful events happened during the week for some participants, which they might refuse to report.

## 5. Conclusions

The present study demonstrates a stable, functional resting-state network of the amygdala, which is connected not only with areas involved in the emotion circuit, but with brain regions associated with high cognitive functions that have also been reported as DMN nodes. Based on resting-state FC values obtained at a one-week interval, we conclude the amygdala network is stable in time and across healthy subjects with low to moderate anxiety; however, this reproducibility is asymmetrical for the right and left amygdala connections. The right amygdala had more significant connections with other brain regions and its FC values were more stable across the week than the left amygdala FC. These results support a hypothesis of functional lateralization of the amygdala. The left amygdala is more responsible for the conscious processing of threats, which produces more variable FC in a resting state, while the right amygdala FC is more persistent due to its greater engagement in continuous automatic evaluation of stimuli. Moreover, functional connections of ROIs taken from the resting-state amygdala network were more stable and prominent in the right hemisphere; in particular, IFGorb and IFGtriang were functionally connected with the largest number of other brain areas. Finally, reproducible values of the amygdala’s FC may provide an additional baseline for further research, as a reference for comparison with experimental and clinical groups. However, lateral asymmetry of the amygdala’s functional connections during rest in healthy subjects may also raise some additional questions regarding changes in FC data in emotion-induced tasks.

## 6. Compliance with Ethical Standards

### Funding

This study was supported by the Russian Scientific Foundation (Grant № 16-15-00300).

### Ethical approval

All procedures performed in studies involving human participants were in accordance with the ethical standards of the institutional and/or national research committee and with the 1964 Helsinki declaration and its later amendments or comparable ethical standards.

### Informed consent

Informed consent was obtained from all individual participants included in the study.

## Abbreviations

ACC: anterior cingulate cortex
AMYG: amygdala
BDI: Beck depression inventory
Caud: caudate nuclei
FC: functional connectivity
fMRI: functional magnetic resonance imaging
HIP: hippocampus,
IFGorb: inferior frontal gyrus, pars orbitalis
IFGtriang: inferior frontal gyrus, triangular part
INS: insula
MTG: middle temporal gyrus
PCC: posterior cingulate cortex
PHG: parahippocampal gyrus
preCG: precentral gyrus
PTSD: post-traumatic stress disorder
ROI: region of interest
SFG: superior frontal gyrus, SFGmed – superior frontal gyrus, medial
STAI: State-Trait Anxiety Inventory
STP: superior temporal pole
UNC: uncus
vlPFC: ventrolateral prefrontal cortex
vmPFC: ventromedial prefrontal cortex

